# Structural and evolutionary features of red algal UV sex chromosomes

**DOI:** 10.1101/2024.12.05.626989

**Authors:** A.P Lipinska, G. Cossard, P. Epperlein, T. Woertwein, C. Molinier, O. Godfroy, S. Carli, L. Ayres-Ostrock, E Lavaut, F. Marchi, S. Mauger, C. Destombe, M.C. Oliveira, E.M. Plastino, S.A. Krueger-Hadfield, M.L. Guillemin, M. Valero, S.M. Coelho

## Abstract

**Background:** Sex chromosomes in red algae have remained relatively understudied, despite their fundamental role in understanding the evolution of sex determination across eukaryotes. In this study, we investigate the structure, gene composition, and evolutionary history of the U and V sex chromosomes in four *Gracilaria* species, which diverged approximately 100 million years ago (MYA).

**Results:** Our findings reveal that UV sex chromosomes, previously identified in green and brown algae as well as bryophytes, have also evolved in red algae, contributing to the diversity of sex determination systems across eukaryotes. The shared orthology of conserved sex-determining region (SDR) genes between *Gracilaria* and distantly related red algae suggests that this system may have originated approximately 390 MYA, making it one of the oldest known sex chromosome systems. The SDR in *Gracilaria* is relatively small but contains conserved gametologs and V-specific genes involved in transcriptional regulation and signaling, suggesting their essential role in sexual differentiation. Unlike the conserved V-specific genes, U-specific genes appear absent, pointing to a dominant role of the V chromosome in sex determination. The evolution of *Gracilaria* sex chromosomes involved recombination suppression, gene relocations, duplications, and potential gene loss. Despite their ancient origin, the sex chromosomes show low levels of degeneration, likely due to haploid purifying selection during the gametophytic phase of the life cycle.

**Conclusion:** This study provides the first detailed characterization of the U and V sex chromosomes in red algae, preparing the ground for future studies on reproductive life cycles and speciation in this understudied group of eukaryotes.

## Background

Most eukaryotic organisms reproduce sexually, but the mechanisms of sex determination and the nature of their sexual systems can differ significantly, even among closely related species [1,2]. Across eukaryotes, sex determination systems can be environmental, epigenetic, or genetic [3] and have been extensively studied in animals, plants, and more recently in brown algae [4–8]. However, red algae (Rhodophyta) have largely been absent from these analyses. Yet, these photosynthetic eukaryotes are a key component of marine ecosystems and among the earliest-diverging lineages to evolve complex multicellularity [9,10]. Despite their ecological, evolutionary, and economic significance, they remain relatively understudied [11]. As a sister clade to Viridiplantae (green algae and embryophytes), red algae offer valuable insights into early eukaryotic evolution [12].

The Florideophyceae is the largest class of red algae, accounting for over 94% of all currently described species [13] (Figure 1A). A unique feature of most Florideophytes is their complex life cycle, which includes three phases: the diploid tetrasporophyte, the diploid carposporophyte, and haploid gametophytes (Figure 1B). Most marine species are thought to be dioicous (i.e., have separate male and female gametophytes) [14], but freshwater red macroalgae display variation in monoicy, dioicy, and trioicy (male, female, and hermaphroditic gametophytes in the same population; see review in [15]) as well as different life cycles (see [16]). In dioicous species, male gametophytes release non-motile spermatia (male gamete), as red algae do not have flagella, while female gametophytes produce a carpogonium (female gamete) that remains on the female thallus [17,18]. Notably, red algae lack a dedicated germline, so any cell within the thallus can differentiate to produce reproductive structures. After fertilization, the carposporophyte develops on the female gametophytic thallus whereby the zygote is mitotically amplified, producing diploid carpospores. The non-motile carpospores are released and form the next phase, the diploid tetrasporophytes. Tetrasporophytes undergo meiosis, generating non-motile, haploid tetraspores that germinate into either male or female gametophytes thereby completing the life cycle (Figure 1B). Interestingly, these characteristics of the life cycle (non-flagellated male gametes and the carposporophyte - a diploid structure, protected and nourished by the parental female haploid gametophyte) suggest that sexual selection can occur in red macroalgae [19]. van der Meer and Todd [20] have shown that the sex determination is controlled by one mendelian locus. The discovery of sex-linked genomic regions in *Gracilaria* spp. [21–23], *Bostrychia moritziana* [24], and *Pyropia haitanensis* [25] further supports that sex determination in this group, is genetically controlled and involves UV sex chromosomes, as observed in bryophytes, some green algal lineages, and brown algae [26]. However, genomic studies on red algal sex determination systems remain limited.

**Figure 1.**
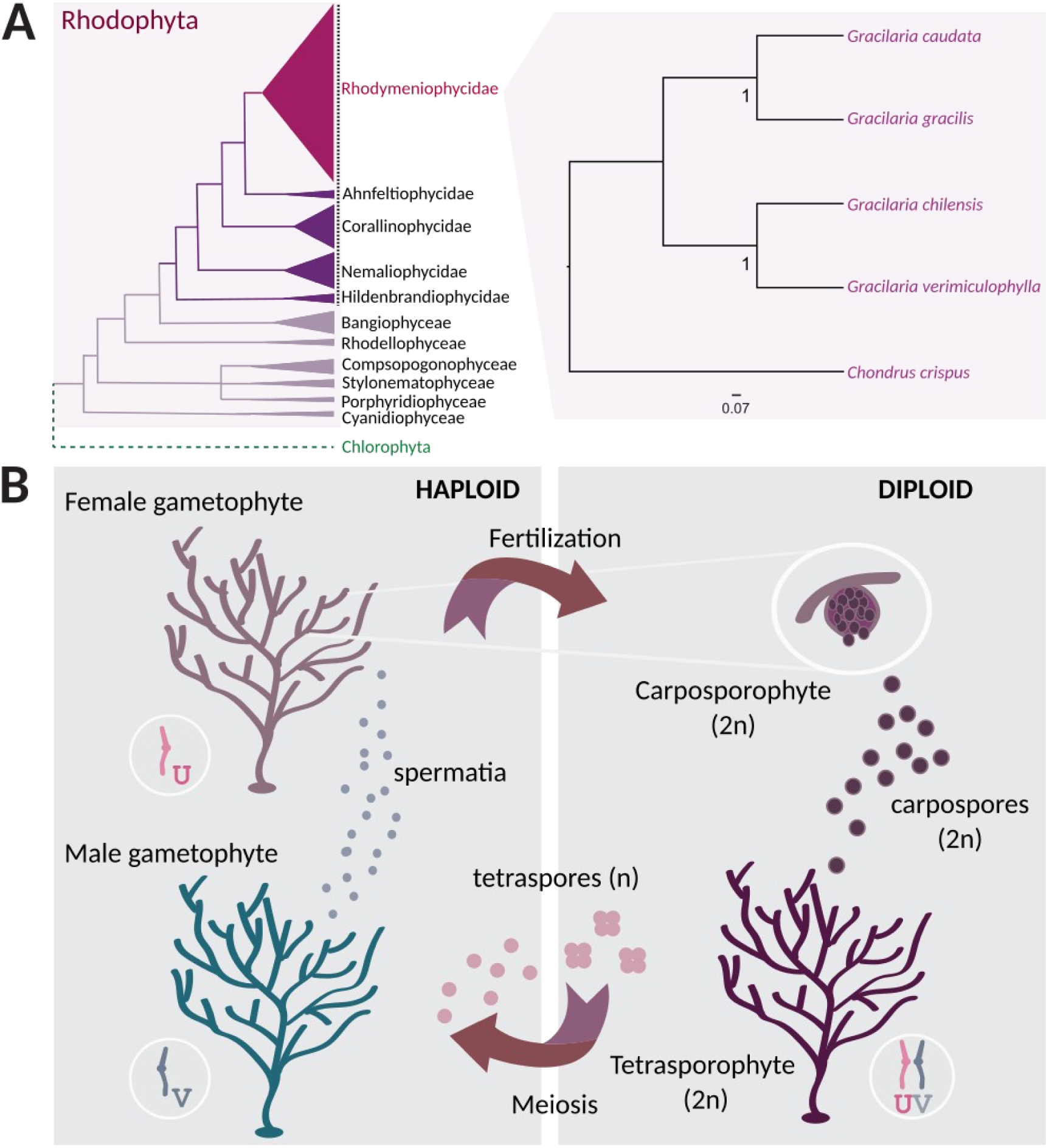
A) Phylogeny of the Rhodophyta and the four *Gracilaria* species used in this study (based on [28]); a dotted line marks the Florideophyceae class. B) Life cycle of *Gracilaria* spp. with alternation between haploid and diploid phases. Haploid male gametophytes produce spermatia which will fertilize female carpogonia giving rise to a diploid carposporophyte that develops on the maternal female thallus. Once mature, carposporophytes will release (diploid) carpospores that will germinate into diploid tetrasporophytes. Tetrasporophytes carry both the U and the V sex chromosomes which are passed on to the haploid spores after meiosis. Haploid tetraspores carrying the V sex chromosome develop into a male gametophyte whereas spores carrying U sex chromosome will produce a female gametophyte. The size of the propagules are different in *G gracilis*: male gametes: 4.5 µm, diploid carpospores 28 µm, haploid spores 15 µm [29].

In this study, we provide the first comprehensive characterization of the UV sex chromosomes in red algae focusing on four species of *Gracilaria* (class Florideophyceae, subclass Rhodymeniophycidae). We explore the structure, gene content, and evolutionary history of *Gracilaria* UV sex chromosomes, as well as the patterns of sex-biased gene expression potentially regulated by these chromosomes. Our results reveal that the sex-determining regions are small, as predicted by [20], and that female development triggers a cascade of up- and down-regulation of many autosomal genes, supporting the idea that male development requires suppression of the female program [27]. We demonstrate that the UV sex chromosomes have remained remarkably conserved over more than 100 million years, and possibly up to 390 million years. Our study emphasizes the stability of core sex-linked genes and resilience of the sex-determining regions throughout the evolution of the genus *Gracilaria* and, more broadly, sheds new light on the evolutionary dynamics of sex chromosomes across the eukaryotic tree of life.

## Results

To define the male and female haplotypes of the sex locus in *Gracilaria* species, we leveraged male reference genomes along with complementary female genomic and transcriptomic data published recently in [30] (Table S1) and also deposited in the Rhodoexplorer database (https://rhodoexplorer.sb-roscoff.fr/home/). Our approach involved the generation of female genome assemblies for all four *Gracilaria* species (see Table S2 for assembly metrics) and the application of various bioinformatic strategies combined with experimental validation, as outlined in the Materials and Methods section. Notably, the male genomes of *G. vermiculophylla*, *G. chilensis,* and *G. gracilis* as well as the female genome of *G. gracilis* represent continuous and high-quality assemblies, whereas the female genomes of *G. caudata, G. chilensis, G. vermiculophylla,* along with the male genome of *G. caudata,* are draft assemblies. We assume that the combination of approaches used here has provided a near-exhaustive list of male and female sex-linked genes and genomic regions in all species (Table S3). It is nevertheless possible that some scaffolds, particularly those that are highly repetitive, may have been missed in the more fragmented assemblies of *G. caudata* male and female genomes or *G. vermiculophylla* and *G. chilensis* female genomes (Table S2) [30]. Therefore, for the sex chromosome structural analysis, including gene density, repeat content, and GC content, we focused only on the most continuous assemblies of male *G. vermiculophylla* and *G. chilensis* along with both the male and female *G. gracilis.* All other analyses concerning sex-linked genes (gene structure, function, expression, and evolution) were conducted across all species and both sexes.

### Sex Chromosome Architecture in Gracilariales

Similar to other organisms with haploid-diploid life cycles, such as those from green algae, bryophyte, and brown algal lineages [26], *Gracilaria* species have haploid sex determination with UV sex chromosomes. In these species, the V chromosome carries the male sex-determining region (V-SDR), while the corresponding U-SDR determines sex in females. To perform a fine-scale characterization of the sex chromosome in the Gracilariales, we used the male chromosome-level assembly of *G. vermiculophylla* [30,31]. *G. vermiculophylla* has 24 chromosomes and 95% of the assembled sequences could be placed in the 24 largest scaffolds, with the male-specific V-SDR located on scaffold 41 (Figure 2A). Consequently, the V sex chromosome of *G. vermiculophylla* has a total size of 5.14 Mbp, featuring a small (0.873 Mbp) central sex-determining region (SDR) bordered by two pseudoautosomal regions (PARs) (Figure 2B). Similarly to *G. vermiculophylla*, the SDRs in the more continuous assemblies of *G. gracilis* male, *G. gracilis* female, and *G. chilensis* male ranged from 0.645-0.920 Mbp (Table 1; Table S3). It appears, therefore, that the size of the sex locus in all examined *Gracilaria* species is relatively small and constitutes less than 2% of the total genome sequence [30,31]. Importantly, we validated previously identified sex markers [21–23] using bulk-segregant analysis and hypervariable genetic markers such as Random Amplified Polymorphic DNA (RAPD), Sequence Characterized Amplified Region (SCAR), and SNPs. These rapid methods proved highly effective in detecting sex markers, even when the SDRs are small.

**Figure 2.**
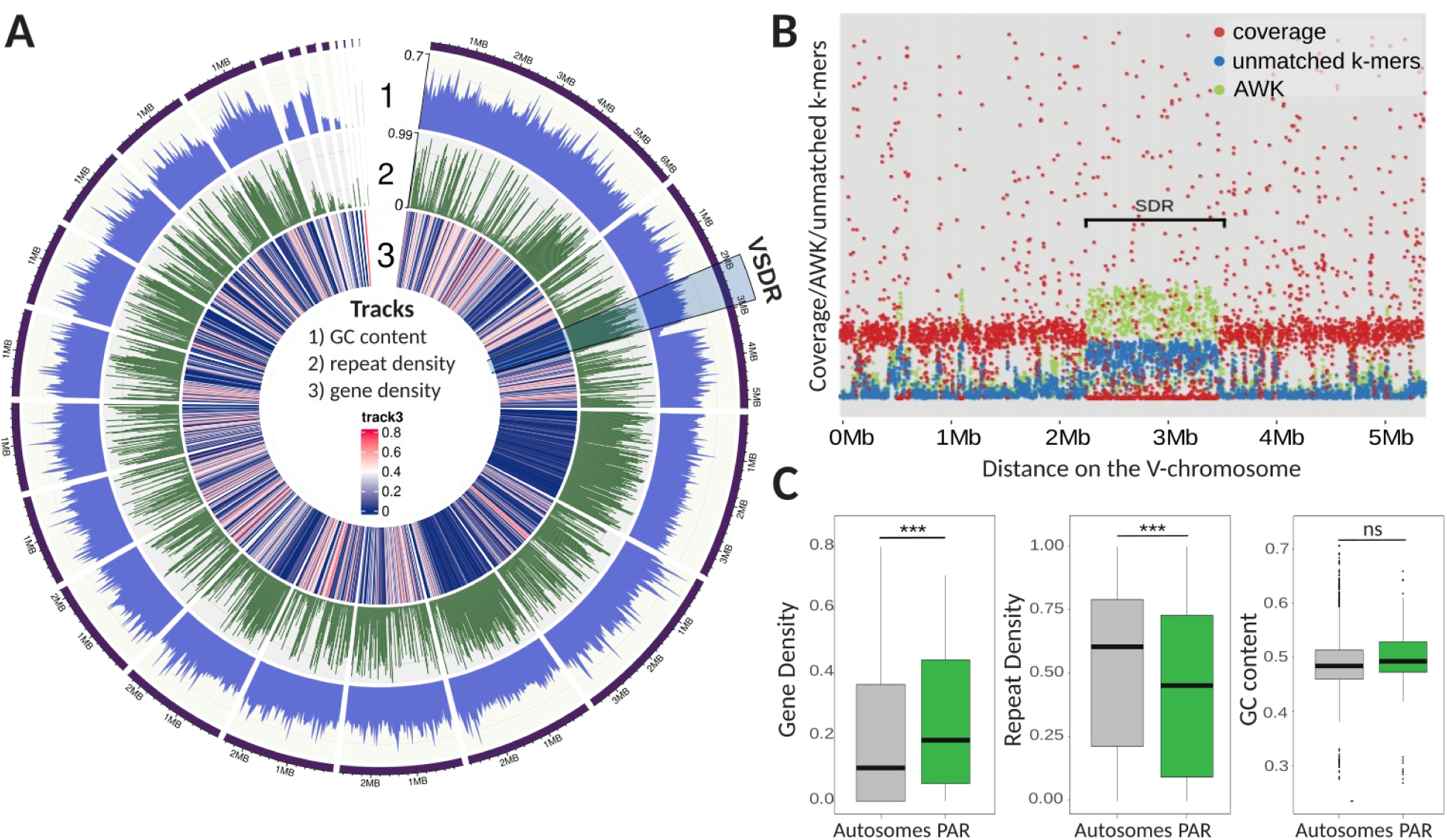
Sex chromosome in *G. vermiculophylla*. A) Genome plot showing GC content, repeat density, and gene density on different chromosomes. V-SDR on Scaffold 41 is highlighted by the blue box. B) Scatterplot of KQ, YGS and coverage of female reads over 1kb-long windows along the scaffold_41 (V-chromosome) in *G. vermiculophylla*. The region spanning from 2,108kb to 2,981kb, displayed no female coverage (red), high AWK values (green dots) and percentage of unmatched k-mers (from KQ and YGS analysis respectively), was retained as the non-recombining sex-determining region (SDR) of the V-chromosome. Green: AWK values, Red: female reads coverage, blue: % unmatched k-mers (YGS, KQ). C) Structural characteristics (gene density, repeat content, GC content) of pseudoautosomal regions (PAR) compared to autosomes. Stars above the boxplots indicate significant differences (***p-value < 0.002, permutation tests, 10k permutations).

**Table 1.**
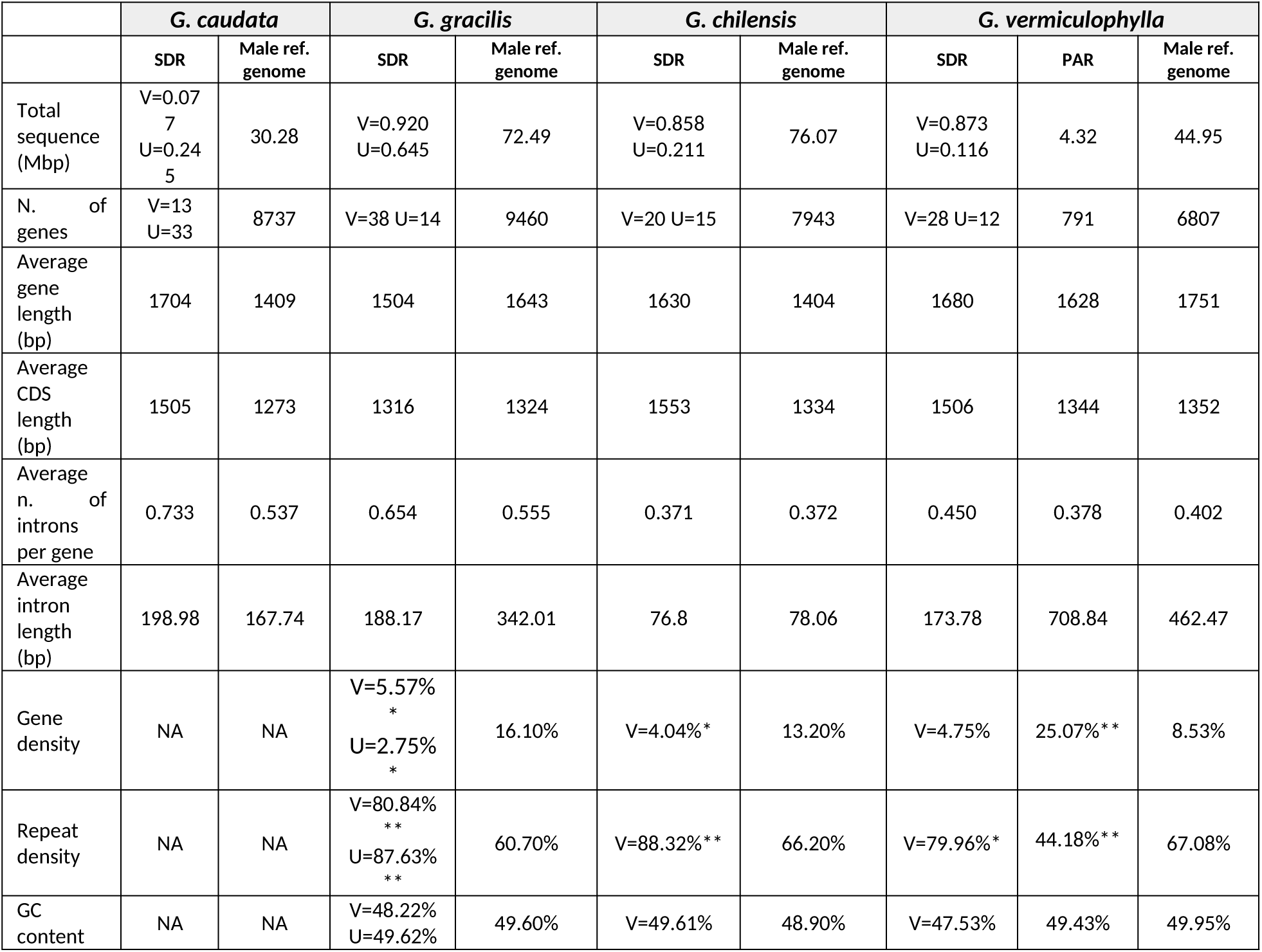
Statistics for several features of the male and female sex-linked scaffolds compared with the complete genome across the four *Gracilaria* species. Asterisk indicates significant difference between the SDR and the autosomes (permutation test of the difference in the mean, *p<0.05, **p<0.001, 10k permutations).

|  | *G. caudata* |  | *G. gracilis* |  | *G. chilensis* |  | *G. vermiculophylla* |  |  |
| --- | --- | --- | --- | --- | --- | --- | --- | --- | --- |
|  | SDR | Male ref. genome | SDR | Male ref. genome | SDR | Male ref. genome | SDR | PAR | Male ref. genome |
| Total sequence (Mbp) | V=0.077<br>U=0.245 | 30.28 | V=0.920<br>U=0.645 | 72.49 | V=0.858<br>U=0.211 | 76.07 | V=0.873<br>U=0.116 | 4.32 | 44.95 |
| N. of genes | V=13<br>U=33 | 8737 | V=38 U=14 | 9460 | V=20 U=15 | 7943 | V=28 U=12 | 791 | 6807 |
| Average gene length (bp) | 1704 | 1409 | 1504 | 1643 | 1630 | 1404 | 1680 | 1628 | 1751 |
| Average CDS length (bp) | 1505 | 1273 | 1316 | 1324 | 1553 | 1334 | 1506 | 1344 | 1352 |
| Average n. of introns per gene | 0.733 | 0.537 | 0.654 | 0.555 | 0.371 | 0.372 | 0.450 | 0.378 | 0.402 |
| Average intron length (bp) | 198.98 | 167.74 | 188.17 | 342.01 | 76.8 | 78.06 | 173.78 | 708.84 | 462.47 |
| Gene density | NA | NA | V=5.57%<br>*<br>U=2.75%<br>* | 16.10% | V=4.04%* | 13.20% | V=4.75% | 25.07%** | 8.53% |
| Repeat density | NA | NA | V=80.84%<br>**<br>U=87.63%<br>** | 60.70% | V=88.32%** | 66.20% | V=79.96%* | 44.18%** | 67.08% |
| GC content | NA | NA | V=48.22%<br>U=49.62% | 49.60% | V=49.61% | 48.90% | V=47.53% | 49.43% | 49.95% |

Next, we investigated if the UV system in red algae also exhibits the structural characteristics typically associated with sex-specific, non-recombining regions, such as lower gene content and accumulation of repeats compared to autosomes [5,32–34]. These patterns are attributed to the suppression of recombination across the SDR which can lead to genetic degeneration unless there is a strong selection on gene function to counteract this effect [35–37]. As expected, we found that the gene density on the sex-linked scaffolds in *Gracilaria* species was significantly lower (p-value < 0.05, permutation test of the difference in the mean, 10k permutations) and the repeat content was considerably higher (p-value < 0.05, permutation test of the difference in the mean, 10k permutations) compared to the autosomes (Table 1; Table S4, S5). However, other indicators of genetic degeneration, such as lower GC content [4,38] or shorter CDS length [33], did not show notable differences between sex-linked and non-sex-linked regions (p-value > 0.05, permutation test of the difference in the mean, 10k permutations, Table 1; Table S4). Furthermore, coding sequences of sex-linked genes in all four species did not display a significant under-representation of optimal codons, typically associated with degeneration in non-recombining regions (e.g. [39])(Mann-Whitney rank test, p-value>0.05, Figure S1, Table S5). Therefore, although typical structural characteristics of non-recombining regions, such as low gene content and higher repeat content, are present in the *Gracilaria* SDR scaffolds, we found little evidence of degeneration in the genes located within these regions. In fact, the SDR genes did not present any distinguishing feature from autosomal genes (Table 1, Table S5).

Previous studies in brown algal UV systems have demonstrated that pseudoautosomal regions (PARs) exhibit unique characteristics, such as the accumulation of taxonomically restricted, evolutionarily ‘young’ genes and may act as cradles for *de novo* gene birth [7,40]. We therefore examined the PARs of the *G. vermiculophylla* sex chromosome. We found that the average gene density in the PAR region of *G. vermiculophylla* was significantly higher than that of autosomes (p-value = 0.0015, permutation tests of the difference in the mean, 10k permutations; Figure 2C). However, we found no evidence of accumulation of evolutionary ‘young’ genes (Figure S2A). The PAR was instead slightly depleted in young genes compared to autosomes (Χ2=16.514, p=0.035), which stands in contrast to the patterns observed in brown algal UV sex chromosomes [7].

In addition, unlike other UV systems, PARs in *G. vermiculophylla* exhibited lower repeat content (p-value = 5 x 10^-4^, permutation tests of the difference in the mean, 10k permutations) (Figure 2C) and similar substitution rates (Ks_PAR_ = 0.00590 vs Ks_auto_ = 0.00678; (p-value = 0.7567, permutation tests of the difference in the mean, 10k permutations), Table S6, Figure S2B) compared to autosomes. Altogether, these characteristics may be explained by high recombination rates on the PAR [41], but currently, recombination maps are not available for any red algal species to confirm this hypothesis.

To summarize, our analyses showed that the SDRs of representative *Gracilaria* species are small and present typical structural characteristics of non-recombining regions, such as lower gene density and higher repeat content, but they exhibit little genetic degeneration. Contrary to the sex chromosomes of other UV systems [7], we found no evidence for distinctive features in the red algal PARs compared to autosomes [7].

### Evolutionary history of the sex-determining regions in *Gracilaria*

To study the evolutionary history of the *Gracilaria* sex chromosomes, we first examined the genes present in the male and female SDRs. The male haplotype harbors between 13 (*G. caudata*) and 38 (*G. gracilis*) protein-coding genes, whereas the female haplotype contains between 12 (*G. vermiculophylla*) and 33 (*G. caudata)* genes (Table S5). The sex-linked genes could be further described as ‘sex-limited’, when present only in one haplotype of the sex locus, or ‘gametologs’ when they have homologs in the SDRs of the opposite sex. The presence of the gametologs is consistent with the two SDR haplotypes having evolved from a common ancestral autosomal region. The proportion of gametologs among the total sex-linked genes was relatively small and varied across species, ranging from 6/46 genes in *G. caudata*, 8/52 genes in *G. gracilis*, 8/35 genes in *G. chilensis,* and 22/40 genes in *G. vermiculophylla* (Figure 3A). In contrast, sex-limited genes may represent ancestral gametolog pairs, where one of the homologs was lost by the haplotypic counterpart. Alternatively, they may have been acquired by the U-SDR or V-SDR specifically, sometime after the sex-determining regions stopped recombining. To gain further insight into which scenario is more likely, we analyzed the conservation of sex-linked gene content across the four *Gracilaria* species, as well as the overall homology of these genes across the species.

**Figure 3.**
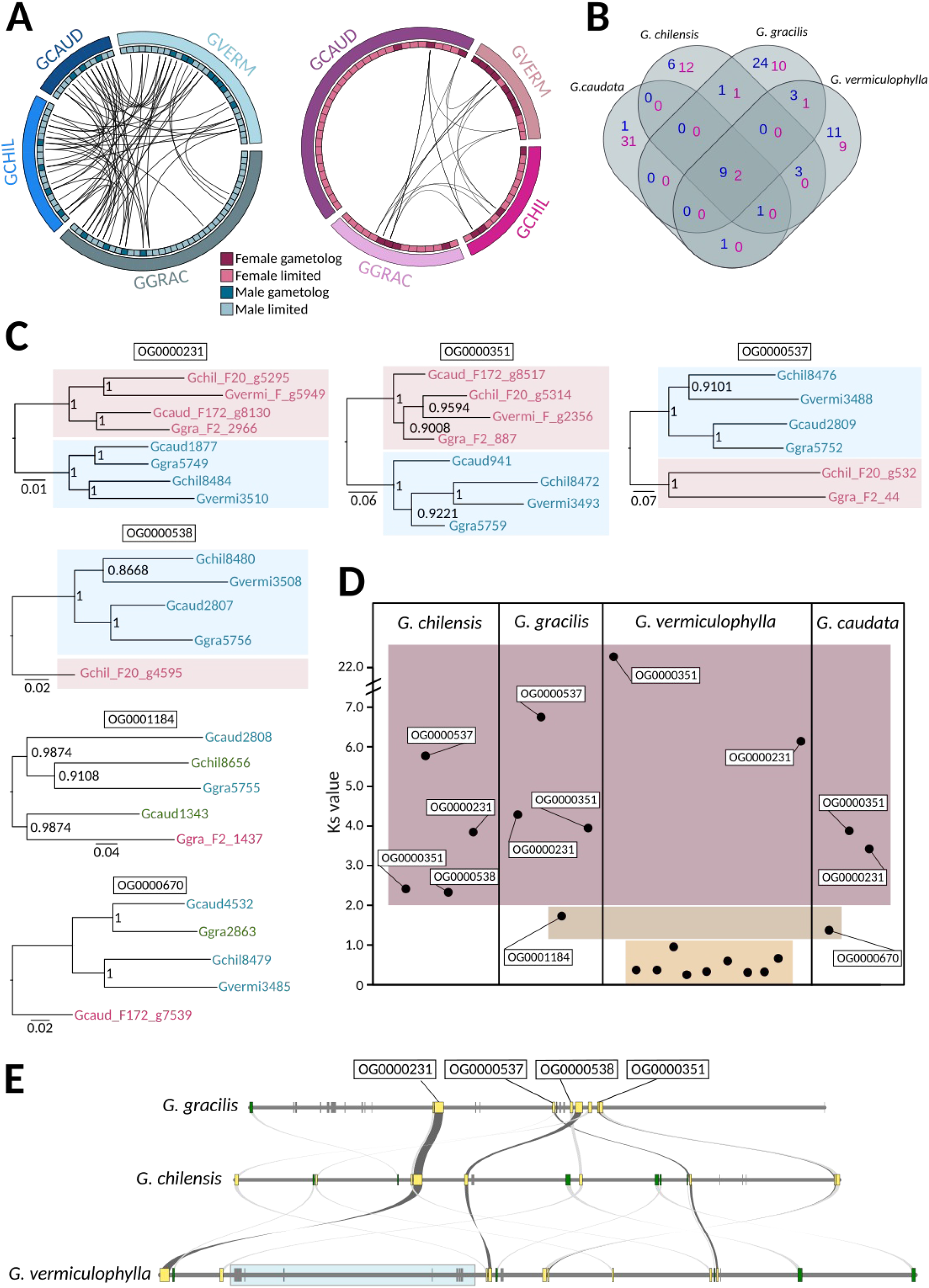
Conservation of SDR genes in the four *Gracilaria* species. A) Schematic representation of the gene content and syntenic relationships among female-linked genes (pink) and male-linked genes (blue) in the four studied species. B) Venn diagram illustrating the orthogroups conservation across the four species (female conserved orthogroups in pink and male conserved orthogroups in blue). See also Table S7. C) Gene trees of conserved gametolog pairs. Bootstrap values (from 1,000 resamplings) are shown next to the branches. D) Synonymous substitutions (Ks) between gametolog pairs per species. Orthogroup names are indicated for gametologs conserved in at least two species. E) Microsynteny plot of the VSDRs in the three species with continuous assemblies. Conserved SDR genes shared by all three species are marked in yellow, and genes shared by two species are marked in green.

We performed an orthology search for protein-coding genes and inferred a total of 9,945 orthogroups (OGs) with 101 of these orthogroups containing at least one sex-linked gene in one of the four species (Table S7). Interestingly, most of the latter orthogroups (80) contained sex-linked genes in only one of the species, and regardless of the evolutionary distance, comparisons resulted in very few shared sex-linked OGs (Figure 3B). This result suggests that the gene content within the *Gracilaria* sex locus is highly dynamic (Figure S3A, Table S8). The similarity between sex-limited SDR genes and their closest autosomal paralog would be consistent with gene duplication events (*i.e*., after the divergence of the U and V) that created either the SDR or the autosomal copy. Given that the expression levels of sex-limited genes (log2(TPM+1)) were significantly lower than those of their autosomal paralogs (paired t-test, t = -2.3133, df = 66, p-value = 0.02383) (Tables S8, Figure S3A), this suggests that the functions of these sex-limited genes may be partially compensated by their autosomal counterparts. This could indicate a potential gradual loss of SDR copies, with the autosomal paralogs arising as functional gene rescues. Alternatively, the SDR copies could be expressed at distinct developmental stages and have specialized roles not captured in our data, as our analysis focuses only on expression in sexual gametophytes. In cases where no autosomal homologs were found, it may point to a species-specific relocation of genes to the SDR or that these could be ancestral SDR genes lost entirely in all the other species (Table S7).

Despite low conservation of SDR gene content, we found two orthogroups with gametolog pairs that were shared across all four species (OG0000231, OG0000351) and two with conserved male gametologs in all species (OG0000537, OG0000538), although the female gametologs have been lost in some of the *Gracilaria* species. Phylogenetic relationships among these conserved gametologs indicate that the SDRs in *Gracilaria* are ancestral and stopped recombining before the speciation event, approximately 100 million years ago [28] (Figure 3C). Remarkably, genes from OG0000231 shared orthology with a male sex-linked gene (Bsm1) identified in another red alga, *Bostrychia moritziana,* from the order Ceramiales [24], which diverged from Gracilariales around 390 MYA [28]. This finding indicates that the sex determining regions in red algae may be even more ancient. Additionally, we found six male-limited genes that were consistently sex-linked in all *Gracilaria* species (Figure 3B, Table S7). These genes may represent the ancestral V-sex chromosome. Notably, no conserved female-limited genes were found on the U chromosome.

Finally, we examined the rates of divergence between gametologs to uncover the ‘evolutionary strata’ of sex chromosomes. By analyzing the synonymous substitution rates (Ks) between gametologs within species, we can estimate the relative timing of successive recombination suppression between U and V sex chromosomes, providing insight into the history and evolutionary dynamics of the SDR regions. Gene-by-gene analysis showed, as expected, that the Ks is highest and saturated for the gametologs associated with the ancestral SDR in all four species (OG000231, OG000351, OG0000538, OG0000537) (Figure 3D, Table S9). Based on gene trees and Ks values for two additional gametolog pairs in *G. caudata* (OG000670) and *G. gracilis* (OG0001184), it appears that these genes became sex-linked at later stages, after which female gametologs were lost, and some male genes relocated outside the V-SDR (Figure 3D). Lastly, the SDR region in *G. vermiculophylla* included a cluster of nine consecutive gametologs (based on the physical position in the male sex chromosome assembly) with lower Ks values (<1), located in an inverted region in relation to the *G. chilensis* SDR (Figure 3D,E). Three of these gametologs share homology with male sex-linked genes in *G. gracilis*, but the phylogenetic analysis suggests independent acquisition, as the genes follow the species tree, with male and female copies grouping together in *G. vermiculophylla* (Figure S4, Table S7). The remaining gametologs are either not sex-linked in any other *Gracilaria* species or lack orthologs altogether, suggesting a recent U-and V-SDR expansion in *G. vermiculophylla* which is further supported by synteny analysis (Figure 3E).

Taken together, our analyses suggest that the sex chromosome system is at least 100 million years old and ancestral to the *Gracilaria* species studied here. The fact that one ancestral gametolog, shared by all four species, is also sex-linked in the distantly related red alga *Bostrychia moritziana*, suggests that this sex chromosome system may have already originated over 390 MYA [24,28]. Although we did not identify clear evolutionary strata on the sex chromosomes, we observed a recent SDR expansion in *G. vermiculophylla* involving a series of inversions and/or gene shuffling events.

### Sex Differences in Gene Expression and Function

We focused on the functions and expression patterns of sex-linked genes, particularly the conserved ones, assuming that these genes play a pivotal role in the cascade of differentiation and development in males and females. Gene ontology analysis revealed that genes in sex-linked scaffolds across all four species were predominantly associated with DNA binding, phosphorylation, and energy metabolism (Table S5, Figure S5). Particularly, conserved male-specific genes included putative key regulators, such as a homeobox transcription factor (OG0003259), a glycosyltransferase (OG0000538), and a serine/threonine-protein kinase (OG0005139). The conserved gametolog pairs were found to be involved in signaling pathways and transcription regulation (e.g., OG0000231: Zinc finger and ankyrin-repeat protein, OG0000351: ATP-NAD kinase, PpnK-type)(Table S5). The significant correlation in transcript abundances between gametolog pairs (log2(TPM+1)) in males and females (Pearson’s r = 0.541, p-value = 0.0248, Figure S3B) suggests that these genes have been retained on the U- and V-SDRs due to their crucial roles during the haploid phase of the life cycle. These roles may be related to sex determination or reproduction but also involve broader gametophytic developmental processes. Therefore, we examined differential expression levels of autosomal genes between male and female gametophytes to identify sex-biased genes that could be regulated by the SDRs.

To explore the extent of sex-biased gene expression, we analyzed transcriptomic data from male and female gametophytes. *Gracilaria* male and female gametophytes are morphologically difficult to distinguish prior to the development of reproductive structures, where sex becomes easily identifiable after the development of spermatangial sori (male reproductive structures) or the carposporophyte (i.e., cystocarp) (Figure 4A). However, in some cases, male gametophytes have been reported to be smaller, as seen in *G. gracilis* and *G. chilensis*, and may exhibit physiological differences, such as variations in growth speed [42–44]. Our analysis included gametophytes representing two developmental stages: younger gametophytes that had just started forming reproductive structures (*G. chilensis* and *G. caudata*) and fully grown mature gametophytes (*G. vermiculophylla* and *G. gracilis*). For *G. chilensis* and *G. caudata*, gametophytes were cultured under controlled laboratory conditions and sex determination was conducted through microscopic observation of male sori or female trichogynes (receptive structures associated with the female gametangia). These gametophytes were young, and because they were grown in isolation, females were fertile but unfertilized and did not develop cystocarps. In contrast, gametophytes of *G. vermiculophylla* and *G. gracilis* were collected directly from the field. These thalli were fully mature, and females were fertilized, bearing cystocarps (which were removed from the tissue prior to RNA extraction).

**Figure 4.**
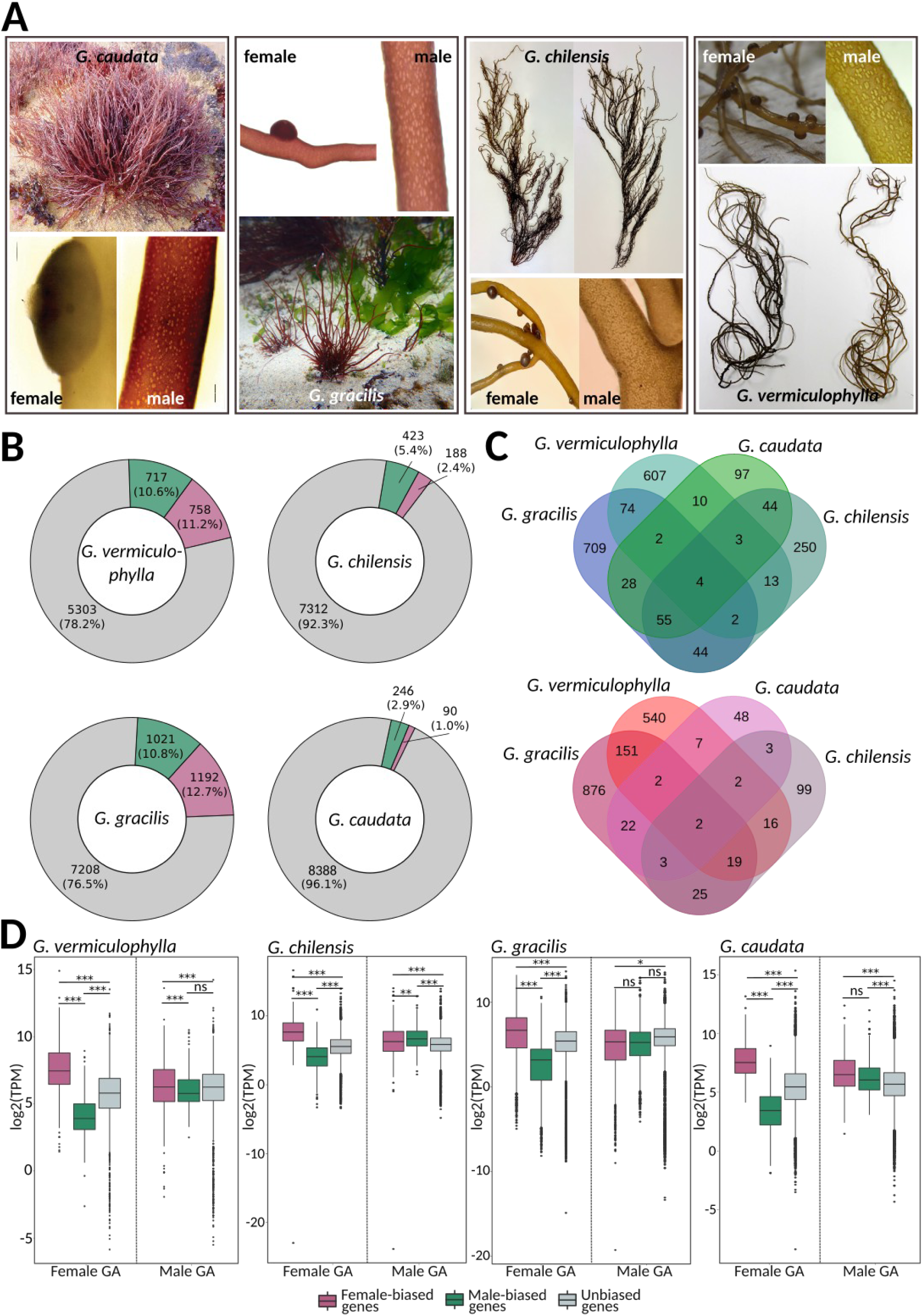
Sex-biased expression in *Gracilaria*. A) Gametophytes of *Gracilaria* showing male gametophytes with the presence of spermatia and female gametophytes with developing cystocarps. B) Proportion of sex-biased genes in *Gracilaria* species, green -male-biased genes, purple - female-biased genes, grey - unbiased genes. C) Venn diagrams representing the number of shared orthogroups (OGs) containing male-biased genes (top) and female-biased genes (bottom) in all four studied species. D) Expression levels (log2(TPM+1) of sex-biased and unbiased genes in male and female gametophytes of *Gracilaria*; green - male-biased genes, purple - female-biased genes, grey - unbiased genes. Significant differences are indicated with asterisk, pairwise Wilcoxon test with Holm correction, ***p-value < 0.0001, **p-value < 0.001, *p-value < 0.05).

We observed significant variability in the proportion of the transcriptome that experiences sex-biased expression patterns depending on the developmental stage of the gametophytes (Figure 4B, Table S10). In *G. caudata* and *G. chilensis*, only a small number of genes were sex-biased (336 and 611 genes, respectively) whereas *G. gracilis* and *G. vermiculophylla* showed moderate levels of biased expression (2,213 and 1,475 genes, respectively). Most sex-biased genes were found in orthogroups unique to each species, indicating that sex-biased expression is highly species-specific. Only four orthogroups exhibited conserved male bias, while two orthogroups showed conserved female bias (Figure 4C). However, we observed a greater number of conserved orthogroups between species pairs representing similar developmental stages: young gametophytes in *G. chilensis* and *G. caudata* versus fully mature and fertilized gametophytes in *G. vermiculophylla* and *G. gracilis*. Functional analysis revealed that male-biased genes of both *G. chilensis* and *G. caudata* were significantly enriched in Gene Ontology (GO) terms related to cell division, DNA replication, and cytoskeletal organization (GO:0044774, GO:0000727, GO:0007020, GO:1905775, GO:0006267, GO:1902975, GO:0006268) (Table S11). These GO terms are characteristic of a highly proliferative tissue, suggesting they could be related to spermatial production in the young male gametophytes. In contrast, female-biased genes of *G. vermiculophylla* and *G. gracilis* female gametophytes were enriched in GO terms involved in response to external stress and defense-related processes (GO:0098869, GO:0045454, GO:0042744) (Table S11). This implies a protective role against environmental challenges, such as intense light exposure, pathogen attacks, or other stressors that trigger reactive oxygen species (ROS) production.

Despite the high turnover in sex biased gene identity, a common pattern emerged across all *Gracilaria* species when we examined the expression levels (log2(TPM+1)) of sex-biased and unbiased genes in males and females. In female gametophytes, female-biased genes (FBGs) were expressed at significantly higher levels, while male-biased genes (MBGs) were expressed at significantly lower levels, compared to unbiased genes. In contrast, male gametophytes exhibited more consistent expression levels across male-biased, female-biased, and unbiased genes, suggesting that sex-biased expression primarily results from the up-regulation or down-regulation of specific genes in females (Figure 4D).

Finally, we examined the evolutionary forces shaping sex-biased genes, focusing on their chromosomal distribution and patterns of coding sequence evolution, measured as non-synonymous to synonymous substitution rates (Kn/Ks). Sex-biased genes are often expected to evolve more rapidly than unbiased genes due to positive selection or relaxed evolutionary constraints [45–48]. Additionally, sex-biased genes might be enriched in PARs if their partial linkage to the SDR provides reproductive or survival advantages, facilitating the spread of advantageous alleles [49]. However, our analysis revealed no significant enrichment of sex-biased genes in the PARs of the *G. vermiculophylla* sex chromosome (Table S12, chi-square test, p-value > 0.05). Furthermore, the evolutionary rates of male- and female-biased genes (measured as Kn/Ks) did not differ from those of unbiased genes, indicating that these genes are likely evolving under purifying selection (Table S13, Figure S6).

In sum, although *Gracilaria* SDRs contain several conserved genes enriched in functions related to signaling and transcription regulation, the downstream networks of sex-biased gene expression are highly variable in this group. Sex-biased genes experience high turnover rates, with little conservation of sex-biased expression among orthologous genes, indicating that such expression is highly species-specific. However, sex bias consistently appears to stem from gene expression regulation in females, suggesting that regulatory elements may play a crucial role in sexual differentiation.

## Discussion

### The UV sex chromosomes of red algae are old

This study provides the first detailed characterization of the U and V sex chromosomes in red algae, focusing on four species of *Gracilaria* that diverged approximately 100 MYA. Our findings reveal that UV sex chromosomes, common among various eukaryotes, such as green algae [50,51], brown algae [7,33], and bryophytes [4,5], have also evolved in this group, adding to the diversity of sex determination systems observed across the eukaryotic tree of life [1,26,52]. The shared orthology of *Gracilaria* conserved SDR gene with a sex-linked gene in more distantly related red algae within the order Ceramiales [24], suggests that this sex chromosome system may have already originated around 390 MYA. This would place it among the most ancient sex chromosome systems, comparable to the UV chromosomes of *Marchantia* and *Ceratodon* [4,5].

### Genomic rearrangements and the evolution of *Gracilaria* sex chromosomes

The sex determining region in *Gracilaria* species is relatively small, yet it harbors a conserved set of gametologs and V-specific genes with functions in transcriptional regulation and signaling, indicating that the ancestral SDR may have evolved to capture genes with crucial roles for sexual differentiation. While a subset of V genes are conservatively sex-linked across red algal species, we found no conserved U-specific genes, suggesting that the V-SDR may play a more crucial role in sex determination. This finding aligns with observations in other *Gracilaria* species, where a rare autosomal recessive mutation (*bi*) enables the development of functional female reproductive structures alongside male spermatangial sori in a male genetic background [27,53]. However, females with the same mutation never develop male reproductive structures indicating that the V chromosome carries a factor essential for male trait expression, while female sexual characters can develop independently of the U chromosome [27,53]. A similar V-dominance has been described in brown algae, where the V chromosome contains a master sex determining gene (*MIN*) [54]. In brown algae, the V-SDR appears to repress the default female program [55,56]. However, in contrast to red algae, the female U chromosome in brown algae is essential for producing functional gametes [54,56,57].

The evolutionary dynamics of the *Gracilaria* SDR appear to involve a series of recombination suppression events, multiple gene relocations and duplications into and/or out of the haploid sex chromosomes, and a potential gene loss. For instance, we observed a recent expansion of the SDR in *G. vermiculophylla*, adding nine new gametologs. Additionally, microsynteny analysis revealed extensive gene shuffling, consistent scrambled gene order between U and V chromosomes, making it difficult to define clear evolutionary strata in specific chromosomal regions [58]. The frequent genomic rearrangements seem to be a defining feature of the UV sex chromosomes and were reported across diverse species groups [4,5,7,51,59,60].

Genomic rearrangements involving sex chromosomes could potentially explain the occurrence of rare sexual phenotypes, known as ‘mixed phases’, observed in tetrasporophytes and gametophytes of various species of red algae in natural populations [20,27,61,62] and in the lab [53,63]. For example, in diploid tetrasporophytes of *Gracilaria gracilis, G. vermiculophylla,* and *G. tikvahiae*, male and/or female reproductive structures have been found next to tetrasporangia. Given that the gametes produced in some of these mixed gametangia were diploid and mitotic recombination seems to occur frequently in red algae [20], it was suggested that mixed diploid phases may result from rare mitotic recombination events between the U and V chromosomes [20,61]. Such events could lead to mosaic individuals with a combination of “normal” cells containing both U and V chromosomes, and cells that are largely homozygous for either U/U or V/V. In this model, sexual characteristics would be independent of ploidy. Heterozygosity at the sex-determining loci, rather than simply being diploid, would drive the tetrasporophytic developmental program. This scenario is supported by our finding that the sex determining regions in *Gracilaria* are relatively small, because mitotic recombination is more plausible if the segregation of sex determining elements rely on a small loci rather than on highly heteromorphic sex chromosomes.

Additional examples of reproductive incongruence include haploid bisexual individuals of *Gracilaria tikvahiae* and *G. caudata* that produced both male and female functional gametangia [27,53]. Crossing experiments between wild-type and bisexual gametophytes followed by segregation analysis of sex phenotypes, revealed that all the bisexual gametophytes were genetically male and carried an autosomal recessive mutation (*bi*). Because gametophytes carrying a female sex chromosome and the *bi* allele developed as females, it was suggested that the mutation affects a locus with a role in the suppression of female program during the differentiation of males [27,53]. Alternatively, the mutant *bi* allele may disrupt or partially inhibit male-specific functions regulated by the V chromosome.

The identification and detailed description of sex chromosomes in this study, combined with advances in genomic data availability for red algae [30,64], will allow the investigation of the features of sex chromosomes and sex-specific differences in gene activity, to better understand origins and stability of ‘mixed-phase’ phenotypes in this important group of eukaryotes.

### Sex chromosomes in *Gracilaria* show low levels of degeneration

Despite the ancient origin of the SDR in *Gracilaria*, we found little evidence of degeneration beyond the accumulation of transposable elements and lower gene density in the sex-determining regions, similar to green and brown algae [7,33,51]. Unlike diploid systems (XY, ZW), where non-recombining regions accumulate deleterious mutations and loose functional genes [65,66], U and V chromosomes are exposed to selection during the haploid phase, slowing degeneration [67]. Aside from the conserved ancestral genes, the SDR content in *Gracilaria* is highly species-specific, with many genes having paralogs on autosomes, suggesting duplication inside/outside the SDR or gene loss. Additionally, we found no significant reduction in codon usage bias among SDR genes, which was reported in other algae [33,51] and mosses [4], again arguing for reduced degeneration levels. These patterns could be explained by a strong selection on the essential functions of U- and V-SDR genes during the haploid phase [68–70]. Another factor that may have limited degeneration is the relatively small size of the SDR and low number of active genes, which could reduce the potential for Hill-Robertson interference among selected sites [65,66,71]. However, given the ancient nature of the SDR, we cannot exclude that degeneration occurred in the past, and what remains are the genes that have been selectively maintained due to their crucial role in sex determination, reproductive processes, or gametophytic vegetative growth. The ancestral gene number on the *Gracilaria* sex chromosome is currently unknown and would require an outgroup without differentiated sex chromosomes, though this is presently unavailable.

The PAR of brown algae show an excess of ‘young’, taxonomically restricted genes, and this feature has led to the hypothesis that the UV sex chromosomes in brown algae may serve as ‘cradles’ for evolutionary novelty [7,40]. A similar pattern was observed in the UV sex chromosomes of some bryophytes, suggesting that this feature may be common among haploid sex chromosomes [7]. However, in *Gracilaria vermiculophylla*, the PAR did not exhibit an enrichment in orphan genes. The gene ‘cradle’ pattern is associated with heterochromatic landscapes, elevated mutation rates, and high transposable element (TE) abundance in brown algal PARs [7]. In contrast, the PAR of *G. vermiculophylla* had significantly lower TE content and similar mutation rates to the autosomes, so it may not provide a favorable environment for the emergence of new genes.

### Regulatory mechanisms of sex-biased gene expression in *Gracilaria*

Conserved SDR genes likely serve as key regulators that orchestrate sex-specific developmental processes, which are essential across different species of *Gracilaria*. However, our findings reveal that sex-biased genes were highly species-specific, with hardly any common sex-biased genes shared across the studied species, a pattern also observed in plants [72] and brown algae [73]. The lack of difference in sequence evolution (Kn/Ks ratio) between sex-biased and unbiased genes implies that sex-biased expression evolves more through regulatory changes rather than through changes in the protein-coding sequences. The consistent pattern where sex-biased expression in females is driven by the upregulation or downregulation of specific genes, while male expression remains constant, points to a conserved regulatory mechanism across species. This is consistent with the observations from bisexual mutants in *Gracilaria*, where expression of the female phenotype was shown to be independent of the U sex chromosome, and the presence of the V chromosome actively suppresses the female program [27,53], which is the opposite of what has been observed in brown algae [56]. Although this is an intriguing pattern, with such phenotypes observed in natural populations at frequencies greater than 1% [61,62,74], we currently lack data on the fine-scale of gene expression across different cell types and developmental stages in red algae.

## Conclusions

In conclusion, this study enhances our understanding of the evolutionary dynamics of sex chromosomes in red algae, demonstrating how these regions have remained remarkably stable despite their ancient origins. The limited evidence of degeneration, coupled with the conserved gene set within the SDR, underscores the importance of these regions in maintaining essential functions related to sex determination and the haploid phase of the life cycle. Our results pave the way for future research to examine changes in sex expression in bisexual haploid and diploid individuals and their putative consequences on the evolution of reproductive life cycles and speciation.

## Material and methods

### Genome assembly and annotation

To define the male and female haplotypes of the sex locus of Gracilariales, we used reference male genome sequences of four *Gracilaria* species complemented with female genomic data published recently in [30], https://rhododev.sb-roscoff.fr/) (see Table S1 for data accession numbers). Female genome assemblies using Illumina 150bp paired-end reads for *G. vermiculophylla*, *G. caudata,* and *G. chilensis* were produced with metaSPAdes v3.12.0 (k=127) [75] and bacterial contigs were purged from the final assembly using BlobTools [76]. Female genome assembly of *G. gracilis* was generated with PacBio long reads using CANU [77]. The assembly was polished with three iterations of RACON v.1.4.20 [78] and errors were corrected using Illumina reads with PILON v.1.23 software [79]. Each female genome assembly was next masked using RepeatMasker v4.0.9 [80] with Dfam v3.0 database [81] and a customized repeat library produced from concatenated outputs of RepeatScout v1.0.6 [82] and TransposonPSI v1.0.0 [83]. RNA-seq reads were trimmed using Trimmomatic v0.39 (TRAILING:3 SLIDINGWINDOW:4:15 MINLEN:50; [84] and mapped to the reference genome using HISAT2 v2.2.1 [85]. RNA-seq mapping was then used as external evidence to annotate protein-coding genes with BRAKER2 v2.1.6 [86].

### Identification of non-recombining sex-specific regions

Several strategies were used to identify candidate male and female SDR scaffolds as described previously for brown algal UV sex chromosomes [7,33]. In order to identify and develop potential sex-linked markers, we mapped male and female reads onto the genome assemblies of each sex using bowtie2 [87] and retained contigs with same-sex coverage within 25% of the genome-wide median and opposite-sex coverage below 80% of genome-wide median. In addition, we adapted and ran previously developed pipelines based on 15bp-long k-mers of male and female reads: YGS adapted to U-V systems [88] and KQ [89] with both DNAseq and RNAseq reads. Contigs that showed up as sex-specific in at least three of the four methods were retained as potentially sex-linked. In order to experimentally confirm sex-linkage, we developed PCR primers in exonic regions of the candidate sex-linked contigs using Primer3 (optimal Tm at 59°C) and used in silico PCR (ipcress, part of the exonerate package, developed for the Ensembl project) to retain markers that may amplify in only one sex. PCR was performed using three males and three females for *G. chilensis* and *G. caudata*, and on nine males and nine females available for *G. gracilis.* Sequences of the PCR primers used can be found in Table S14. We used previously developed markers by Krueger-Hadfield et al. [23] to identify in silico male and female-specific regions in *G. vermiculophylla*. Together, these methods allowed us to confidently identify scaffolds corresponding to the male (V) and female (U) SDR haplotypes in each species (Table S3).

### Orthology inference and evolutionary analysis

We inferred homologous genes between male and female gametophytes of each species using reciprocal blast of protein sequences (e-value = 1×10^-7^, Smith-Waterman algorithm). Pairwise orthologs with both V-linked and U-linked genes are referred to as gametologs. We aligned orthologous gene coding sequences, using the codon-based approach in translatorX v1.1 [90] that implements MAFFT alignment method and Gblocks v0.91b [91,92]. We computed synonymous (Ks) and non-synonymous (Kn) substitution rates of aligned pairwise orthologs using the maximum-likelihood inference of CodeML program in PAML v4.9 [93].

We inferred orthology of protein-coding genes across the four *Gracilaria* species using Orthofinder v2.5.2 [94] with default parameters. We aligned single-copy orthologs (SCO) i.e., orthogroups with a single gene in each of the four species, following the same method described above for pairwise orthologs between males and females. We computed ω (Kn/Ks) using ‘model M0’ in codeml, PAML v4.9 [93].

### GO term analysis

Predicted genes and OGs were blasted against the UNIREF90 non-redundant protein database [95] with blast (v.2.9.0). Annotation was performed using Blast2GO [96], as well as the InterProScan [97,98] prediction of putative conserved protein domains. Gene set enrichment analysis of U-SDR and V-SDR genes were carried out separately for each gene set per species using Fisher’s exact test implemented in the TopGO package [99], with the weight01 algorithm. We investigated enrichment in terms of molecular function and biological process ontology and reported significant GO terms with p-value < 0.05, after Benjamini-Hochberg correction for multiple testing.

### Phylogenetic trees

Gene trees of orthogroups inferred by Orthofinder [94] and containing at least one pair of gametologs in at least one species were inferred using MrBAYES [100] with the BLOSUM62 substitution model with 100k generations (sampling trees every 100 generations with 1000 initial burnin tree samples). Posterior probabilities of node supported are indicated on the trees.

### Sex biased gene expression analysis

To infer gene expression levels, we used kallisto v.0.44.0 [101] using 31-base-pair-long k-mers and 1000 bootstraps. Transcript abundances were then summed within genes using the tximport v3.19 package in R v.4.3.1 [102] to obtain the expression level for each gene in transcripts per million (TPM). Differential expression analysis was done in DESeq2 v3.19 package [103,104] in R v.4.3.1, applying FC (fold change) ≥ 2 and Padj < 0.05 cut-offs (p-values were adjusted using the Benjamini-Hochberg approach). All samples used in the gene expression analysis can be found in Table S1.

## Declarations

### Ethics approval and consent to participate

Not applicable.

### Consent for publication

Not applicable.

### Availability of data and materials

The datasets analyzed during the current study are available in the NCBI repository (see Table S1 for accession numbers) and were included in this published article [30]. Genome assemblies and annotations were additionally deposited on the Rhodoexplorer platform (https://rhodoexplorer.sb-roscoff.fr/).

## Funding

This project was supported by start-up funds from the College of Arts and Sciences at the University of Alabama at Birmingham to S.A.K.-H.; ANID NCN2024-037 and FONDECYT 1221456 and 1221477 to M.L.G.; the International Research Networks DEBMA “Diversity, Evolution and Biotechnology of Marine Algae” (CNRS GDRI 0803) and DABMA “Diversity, Adaptation, and Biotechnology of Marine Algae” (CNRS IRN 00022); the ERC (grant number 864038 to S.M.C.); the ANR project IDEALG (ANR-10-BTBR-04, “Investissements d’Avenir, Biotechnologies-Bioressources”); core funding was provided by the CNRS and Sorbonne University to the International Research Laboratory France-Chile, Evolutionary Biology and Ecology of Algae, CNPq (304776/2022-0 to MCO; 308713/2021-4 to EMP), and an NSF CAREER award (DEB-2141971/DEB-2436117 to S.A.K.-H.)

### Authors’ contributions

*Produced datasets and acquired funding: MLG, SAKH, MV, SMC, MCO; Performed experiments: APL, OG, SM, LA, SC, CD, GC, SAKH, EL*

*Analyzed datasets:: APL, GC, PE, TW, CM, SMC; Interpreted data: APL, SMC*

*Writing-Original draft: APL, GC, SMC. Writing-Review and Editing: APL, SMC, MLG, MV, CD, SAKH. All authors read and approved the manuscript.*

## Acknowledgements

We are grateful to the Roscoff Bioinformatics platform ABiMS (http://abims.sb-roscoff.fr), part of the Institut Français de Bioinformatique (ANR-11-INBS-0013) and BioGenouest network, and the Max Planck Institute for Biology Tubingen for providing computational resources. We thank Josue Barrera-Redondo and Hollie Putnam for helpful discussions. We also thank K. Hill-Spanik for help with *Gracilaria vermiculophylla* collections.

## Abbreviations

SDR: sex determining region
PAR: pseudoautosomal region
FBG: female-biased gene
MBG: male-biased gene
TE: transposable element
MYA: million years ago
OG: orthology group

## Supplementary Materials

Table S1. Data used in this study, published in (Lipinska et al. 2023).

Table S2. Gracilaria female genomes assembly statistics.

Table S3. Sex-linked scaffolds/genomic regions per species.

Table S4. Repeat content, coding sequence and GC content in sex-linked scaffolds. Table S5. Characteristics of sex-linked genes.

Table S6. Synonymous (Ks) and non-synonymous (Kn) divergence between PAR and autosomal genes in *G. vermiculophylla* (PAML4, codeml).

Table S7. Gene orthology analysis across the four Gracilaria species, highlighting the orthogroups containing sex linked genes (blue - V-SDR genes, red - U-SDR genes).

Table S8. Expression of SDR genes and their autosomal paralogs (log2(TPM+1).

Table S9. Synonymous (Ks) and non-synonymous (Kn) divergence between gametologs per species (PAML4, codeml).

Table S10. Sex biased gene expression (transcriptos per million, TPM) in *Gracilaria* species. Table S11. Functional enrichment analysis of GO-terms among sex-biased genes.

Table S12. Distribution of sex-biased genes on the chromosomes of *G. vermiculophylla*. Table S13. Kn/Ks analysis of sex-biased and unbiased genes in *Gracilaria* species.

Table S14. PCR primers used for experimental validation of sex linkage.

## Supplemental Figures Legends

**Figure S1.**
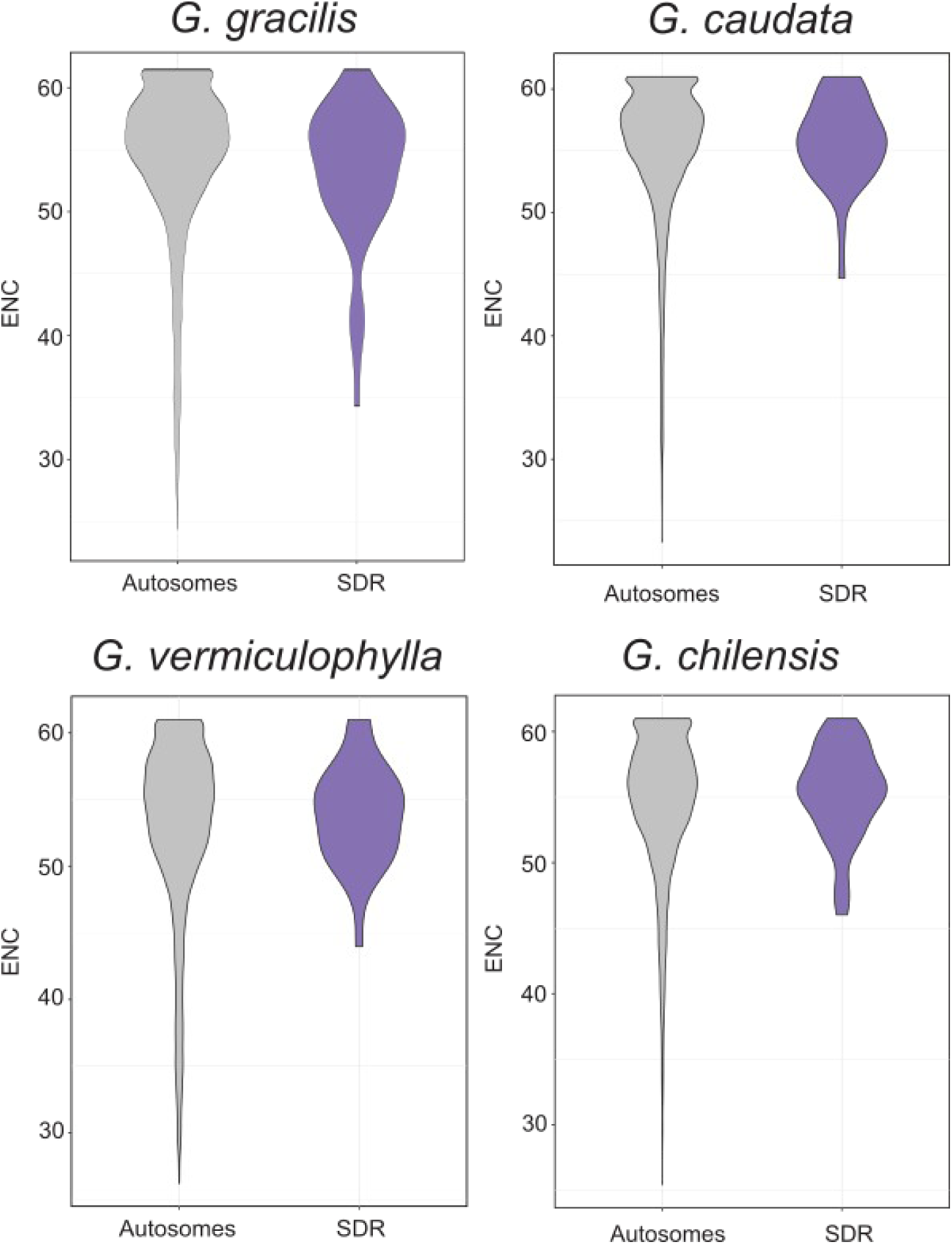
Codon usage bias (ENC’) values for autosomal and sex-linked genes across four *Gracilaria* species. The analysis revealed no significant differences between gene groups (Mann-Whitney U test, p-value > 0.05).

**Figure S2.**
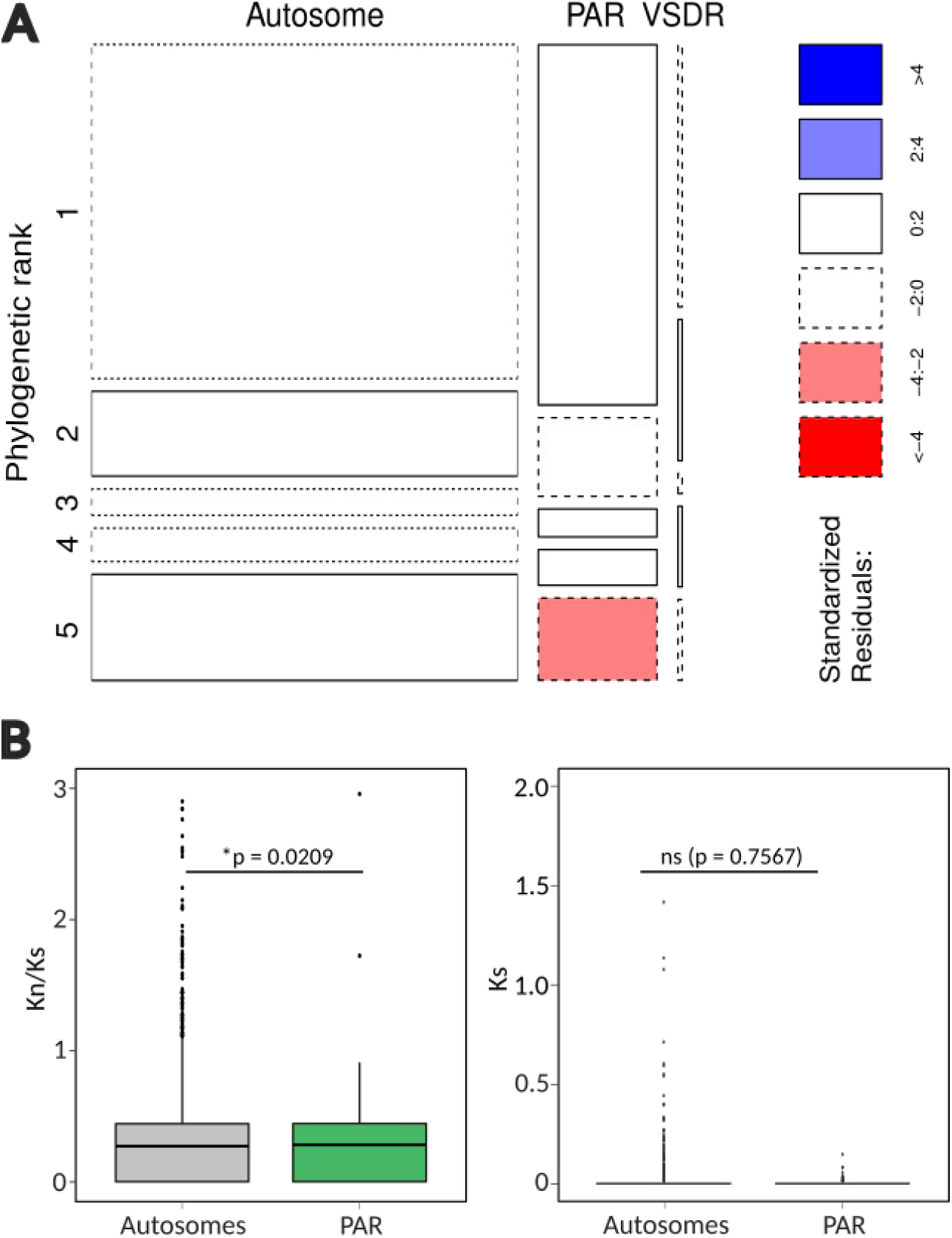
Characteristics of the pseudoautosomal regions (PARs) compared to autosomes in *G. vermiculophylla*. A) Mosaic plot showing that the family-level (rank 5) genes are depleted in the sex chromosome. Phylogenetic ranks (1 - cellular organisms, 2 - Eukaryota, 3 - Rhodophyta, 4 - Rhodymeniophycidae, 5 - Gracilariaceae). B) Non-synonymous/synonymous (Kn/Ks) and synonymous (Ks) substitution rates of the PAR and autosomal genes (permutation tests of the difference in the mean, 10k permutations).

**Figure S3.**
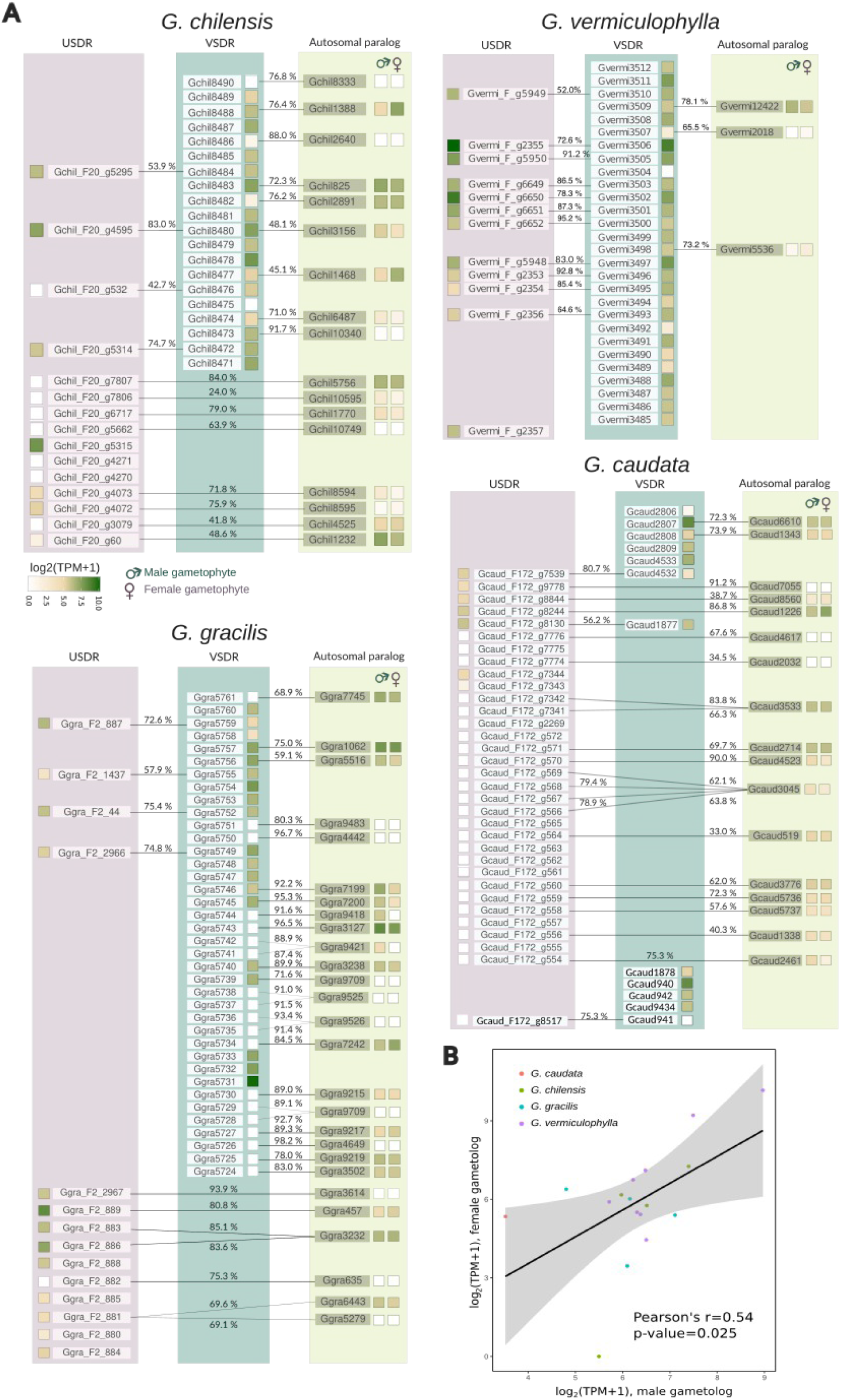
A) Expression levels (log2(TPM+1)) of sex-linked genes (USDR in purple and VSDR in blue-green) and their autosomal paralogs (light green) in the four *Gracilaria* species. Percent of protein sequence identity are indicated above the lines. B) The correlation in transcript abundances between gametolog pairs (log2(TPM+1)) in males and females of the *Gracilaria* species.

**Figure S4.**
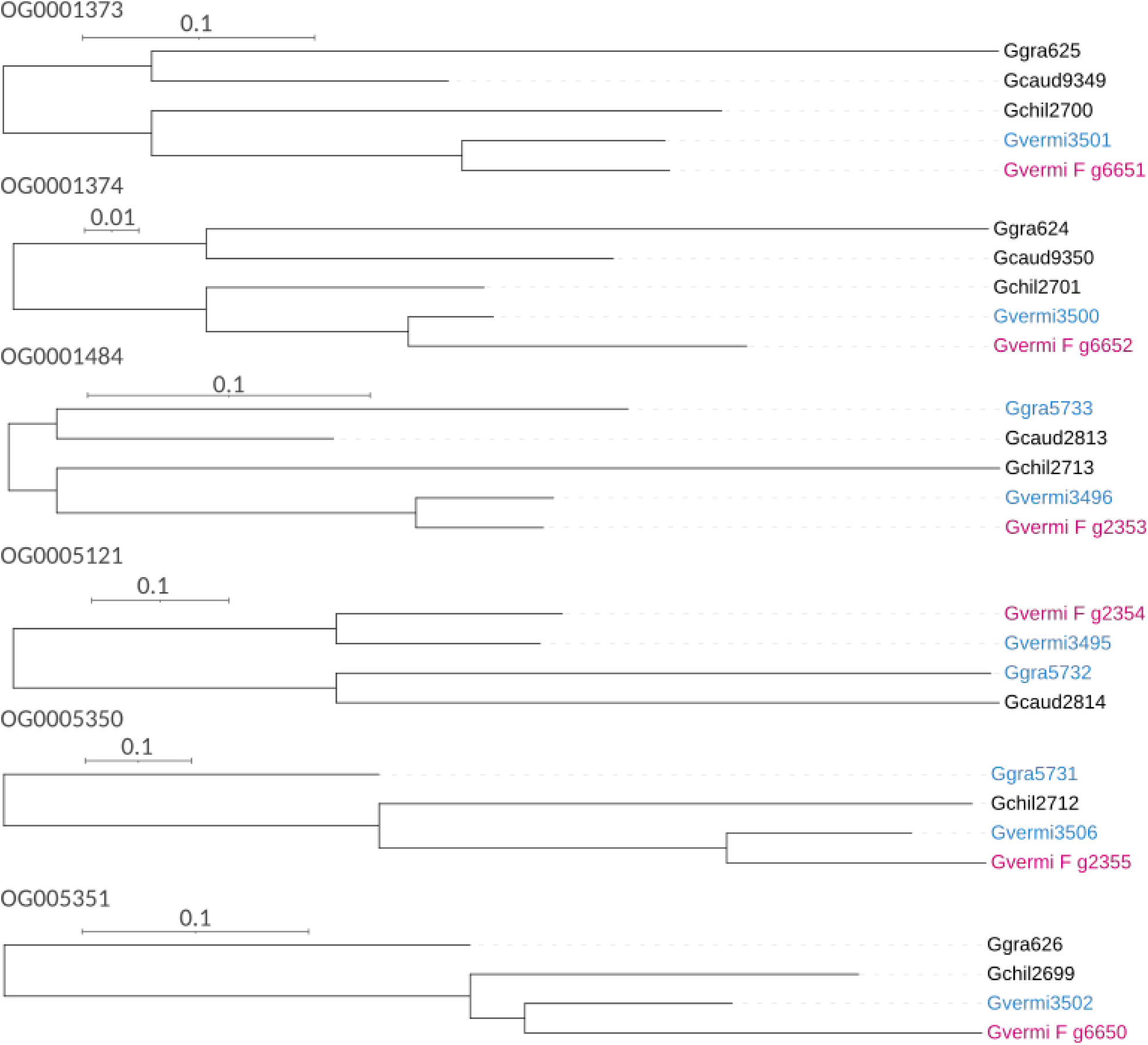
Phylogenetic analysis suggesting the independent acquisition of certain gametolog pairs in *G. vermiculophylla*.

**Figure S5.**
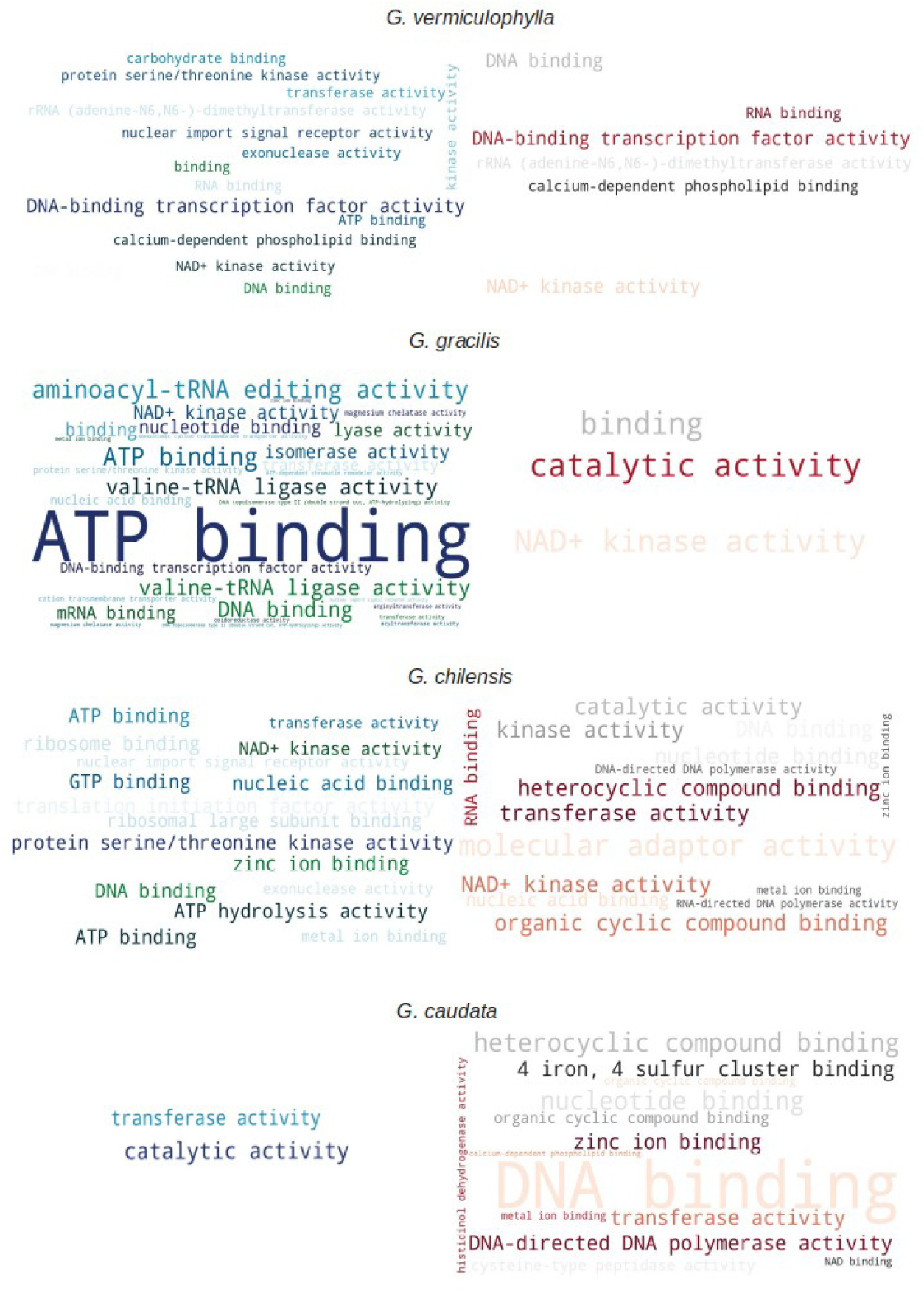
Wordcloud visualization of Gene Ontology (GO) terms enriched in sex-linked genes across all four *Gracilaria* species.

**Figure S6.**
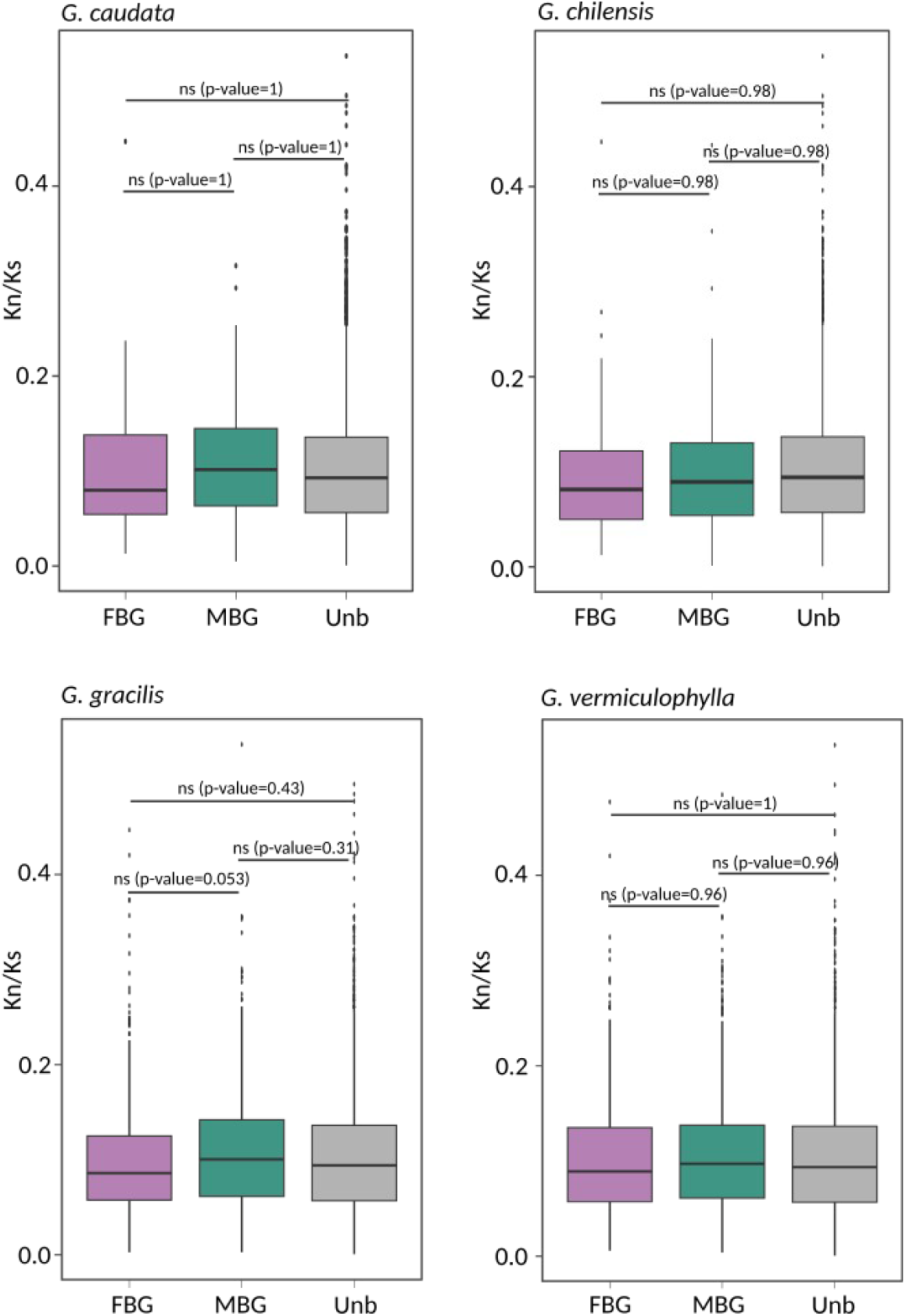
Evolutionary rates (measured as Kn/Ks) of male- and female-biased genes compared to unbiased genes across all four *Gracilaria* species (codeml, model M0, PAML4) showing no statistical difference between the groups (pairwise Wilcoxon test with Holm correction).

